# Novel feature selection method via kernel tensor decomposition for improved multi-omics data analysis

**DOI:** 10.1101/2021.05.21.445049

**Authors:** Y-h. Taguchi, Turki Turki

## Abstract

**Background:** Feature selection of multi-omics data analysis remains challenging owing to the size of omics datasets, comprising approximately 10^2^–10^5^ features. In particular, appropriate methods to weight individual omics datasets are unclear, and the approach adopted has substantial consequences for feature selection. In this study, we extended a recently proposed kernel tensor decomposition (KTD)-based unsupervised feature extraction (FE) method to integrate multi-omics datasets obtained from common samples in a weight-free manner.

**Method:** KTD-based unsupervised FE was reformatted as the collection of kernelized tensors sharing common samples, which was applied to synthetic and real datasets.

**Results:** The proposed advanced KTD-based unsupervised FE method showed comparative performance to that of the previously proposed KTD method, as well as tensor decomposition-based unsupervised FE, but required reduced memory and central processing unit time. Moreover, this advanced KTD method, specifically designed for multi-omics analysis, attributes *P*-values to features, which is rare for existing multi-omics–oriented methods.

**Conclusions:** The sample R code is available at https://github.com/tagtag/MultiR/

## Background

Feature selection with multi-omics datasets has been a long-standing challenge for bioinformatics. Among the numerous proposed methods adapted to multi-omics data analysis [1, 2], only few are capable of performing feature selection. Most of these methods fail to implement feature selection because multi-omics data analysis has a strong tendency to involve a small number (= *n*) of samples with a large number (= *p*) of features, commonly referred to as the *large p small n* problem [3], posing difficulty for accurate feature selection. Features should have a sufficiently small *P*-value to be selected under the null hypothesis. Since the raw *P*-values must be heavily corrected for multiple comparisons when dealing with multi-omics datasets, *P*-values become larger and inevitably less significant; thus, attributing significant *P*-values to individual features, even after correction, is difficult. However, since the number of samples (i.e., conditions) is less than that of features (i.e., variables), labels or values attributed to samples can be accurately predicted by any model (when the number of conditions is less than the number of variables, either the labels or values attributed to samples may be predicted, even if the variables are purely random numbers).

In multi-omics analysis, it is difficult to obtain large sample sizes since multiple observations, each of which corresponds to individual omics approaches, must be performed. In this sense, the required cost and time are multiplied in proportion to the number of omics approaches considered. This often results in a smaller number of samples to which multi-omics measurements are performed when only limited experimental resources are available.

Principal component analysis (PCA) and tensor decomposition (TD)-based unsupervised feature extraction (FE) [4] have been proposed to be applied to feature selection for addressing the *large p small n* problem. Thus, these approaches are also suitable for feature selection in multi-omics analysis. Recently, the TD-based method was extended to kernel TD (KTD)-based unsupervised FE [5], which was applied to integrated analysis of N6-methyladenosine (m6A) and gene expression data [6]. These methods attribute *P*-values to features, which is critical since this enables evaluation of the significance of the selected features, which is rarely possible using other methods applicable to multi-omics datasets [1, 2]. In spite of this advantage, there are limitations of PCA, TD, and KTD-based unsupervised FE when applied to multi-omics data analysis. PCA is inferior to TD when aiming for an integrated analysis of multiomics data. PCA failed to identify genes whose expression and methylation levels are altered simultaneously, but TD could [7]. Although KTD and TD successfully integrated two omics data, they could not achieve the following two points,

1. Reduction of required memory and long CPU time
2. Integration of more than two types of omics data

simultaneously (see Discussions below). We here described a modification of the KTD-based unsupervised FE method to be more suitable for multi-omics data analysis. Although only a small modification was implemented, it nevertheless resulted in more flexibility for multi-omics data analysis, which was verified using synthetic and real data.

Before explaining the results, we briefly discuss the relationship between feature selection and feature extraction, both of which are employed when we are forced to deal with the *large p small n* problem. The former, feature selection, is more straightforward than the latter; it reduces the number of features less than the number of samples. On the other hand, feature extraction is more indirect, since it generates a limited number of new features from the original large number of features. In this study, we have employed a mixed strategy of these two. We first generated new features using feature extraction and selected features using the generated features.

## Methods

### Extended KTD-based unsupervised FE method

Suppose that we have *K* multi-omics datasets with *N*_*k*_ features formatted as tensors sharing sample indices *j*_1_, …, *j*_*m*_ as:

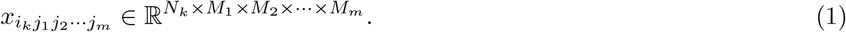

*j*_*s*_, (1 ≤ *j*_*s*_ ≤ *M*_*s*_) refers to the *j*_*s*_th measurement in the *s*th experimental type. *M*_*s*_, (1 ≤ *s* ≤ *m*) is the number of measurements in the *s*th experimental type. Typical examples of *m* experimental conditions include human subjects, tissues, and time points. For example, if the measurements are performed for *M*_2_ tissue types from *M*_1_ individuals at *M*_3_ time points, the total number of samples is *M*_1_×*M*_2_×*M*_3_. *i*_*k*_, (1 ≤ *i*_*k*_ ≤ *N*_*k*_) refers to the *i*_*k*_th feature of the *k*th omics dataset. When *K* types of omics data are measured for each sample, *k* ∈ [1, *K*].

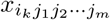 can be kernelized as

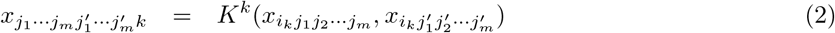

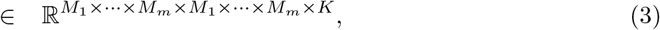

where *K*^*k*^ is an arbitrary kernel applied to 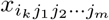. Higher-order singular-value decomposition (HOSVD) [4] is then applied to 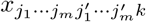, resulting in eq. (4),

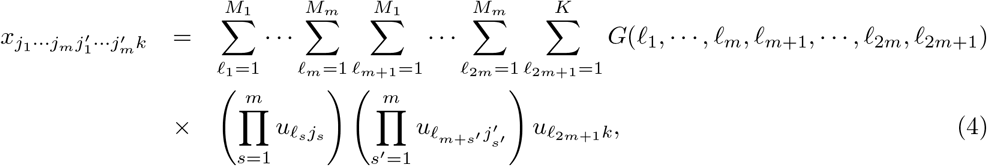

where *£*_*s*_, (1 ≤ *s* ≤ 2*m*) refers to the *£*_*s*_th singular-value vectors attributed to the *s*th experiment type for 1 ≤ *s* ≤ *m* and the (*s* − 2*m*)th experiment type for *m* + 1 ≤ *s* ≤ 2*m*, respectively. *£*_2*m*+1_ refers to the *£*_2*m*+1_th singular-value vector attributed to the omics datasets, 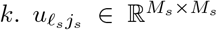 and 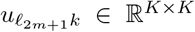 are singular-value matrices, which are also orthogonal matrices, 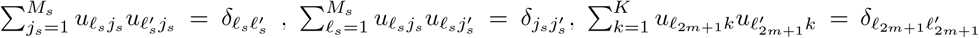, and 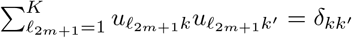 where 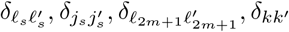 are Kronecker’s delta. Because of symmetry, 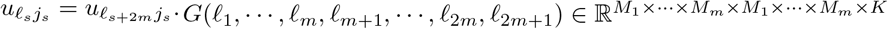 is a core tensor that represents the weight of individual terms composed of the products of singular-value vectors. Here, one should note that 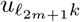 represents the balance (weight) between multi-omics datasets, which usually must be pre-defined manually in the case of a conventional supervised learning approach for FE.

Next, the 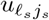 values that are of interest from the biological point of view (e.g., distinct values between the two classes being compared) must be identified. Using these selected 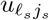 values, singular-value vectors are derived and assigned to *i*_*k*_s as

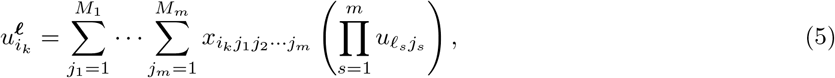

where ***ℓ*** = (*ℓ*_1_, …, *ℓ*_*m*_).

Finally, *P*-values are attributed to *i*_*k*_ assuming that the 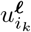 values obey a multivariate Gaussian distribution (null hypothesis) as

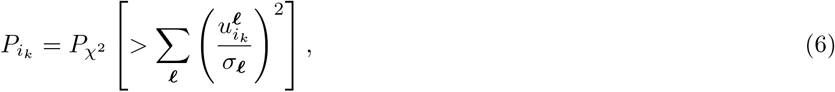

where 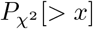 is a cumulative *χ*^*2*^ distribution in which the argument is larger than *x* and *σ*_*ℓ*._ is the standard deviation. Here, summation is taken over *ℓ*_*s*_ selected as being of interest. *P*-values are then computed by the `pchisq` function in R [8].

The obtained 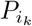 values are corrected using the Benjamini-Hochberg (BH) criterion [4] and the *i*_*k*_ values associated with the adjusted *P*-values less than the established threshold (typically 0.01) are selected. Correction by the BH criterion is performed by the `p.adjust` function in R with the option of method=“BH.”

### Synthetic dataset

A synthetic dataset was derived in the form of a tensor, *x*_*ijk*_ ∈ ℝ^*N* × *M* × *K*^, as

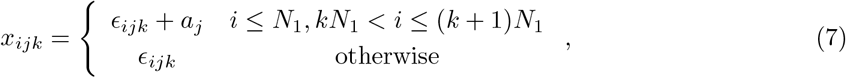

where

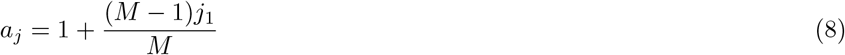

and *ε*_*ijk*_ ∈ (0, 1) is a uniform random real number that emulates the residuals. *N* is the number of variables, *j* is a variable that adds order to the *j*th sample, and *k* is the *k*th omics data. This synthetic dataset assumes that only the top *N*_1_ features among *N* features have dependency on *i* independent of *k. x*_*ijk*_, (*kN*_1_ *< i* ≤ (*k* + 1)*N*_1_) also has dependence on *j*, but in an omics (*k*)-dependent manner. The dependence on *j* is a linear increase upon *j*.

One hundred ensembles of *x*_*ijk*_ were generated and the performances were averaged. First, a linear kernel was generated as

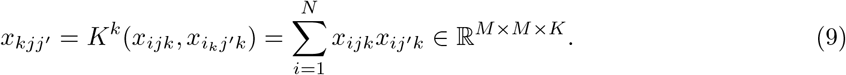

Next, HOSVD was applied, resulting in

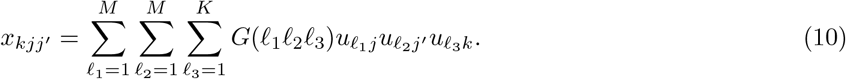

(for the dimensions of the datasets, *N, M, andK*, see the legend to Table 1).Since it was observed that *u*_2*j*_ always had the largest correlation with *a*_*j*_ and *u*_1*k*_ was always constant, regardless of *k, u*_2*j*_ and *u*_1*k*_ were employed to compute

**Table 1.** Confusion matrix when applying KTD-based unsupervised FE to a synthetic dataset (*N* = 1000, *N*_1_ = 10, *M* = 10, *K* = 3).

| | adjusted $P_i \leq 0.01$ | adjusted $P_i > 0.01$ |
| --- | --- | --- |
| $i \leq N_1$ | 7.06 | 2.94 |
| $i > N_1$ | 0.04 | 989.96 |

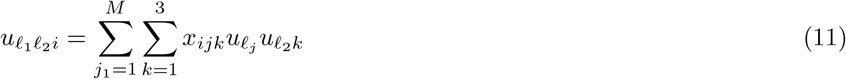

for attributing the *P*-values

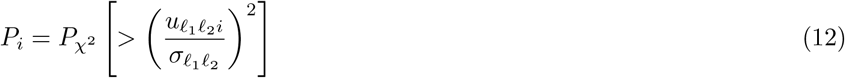

The *P*_*i*_ values were corrected using the BH criterion and the *i* values associated with an adjusted *P*_*i*_ less than 0.01 were selected.

To demonstrate the difficulty of this task, linear regression was used as an alternative method

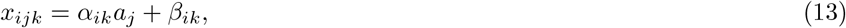

where *α*_*ik*_ and *β*_*ik*_ are regression coefficients, and *a*_*j*_ is defined in eq. (8). Since *x*_*ijk*_s with distinct *k* differ, distinct models were applied to each. After computing BH-corrected *P*-values, the *i*s associated with adjusted *P*-values less than 0.01 were selected.

When the least absolute shrinkage and selection operator (lasso) [9] was applied, the maximum number of features selected were considered, although lasso can select at most as many as *M* features, which is less than the number of features coincident with *a*_*j*_, 2*N*_1_.

When random forest (rf) [10] was applied to synthetic data sets, there were two ways to select features. First, features with nonzero importance were selected. Next, in order to reduce the number of selected features, the top most 2*N*_1_ features having a larger absolute importance were selected.

### Multi-omics hepatitis B virus (HBV) vaccine dataset

The real multi-omics HPV vaccine datasets were based on 75 samples measured in 15 individuals at five subsequent time points (i.e., 0, 1, 3, 7, and 14 days) after HBV vaccine treatment. Gene expression was measured using RNA-sequencing technology and methylation profiles were measured using microarray technology. The proteome was measured for whole blood cells (WBCs) as well as plasma. Since this multi-omics dataset is composed of four types of omics data measured for 15 individuals at five time points, it is thus formatted as a tensor.

Gene expression and methylation profiles were retrieved from the Gene Expression Omnibus (GEO) database using the GSE155198 and GSE161020 datasets, respectively. For gene expression profiles, the GSE155198 RAW.tar dataset was available in the Supplementary File section of GEO. Individual files that included count number of mapped reads toward genes were collected and integrated as a single file. Individual files are named according to the format “GSM*XXXXXXX_*GR*nn_*V*m*.count.txt.gz,” where *XXXXXXX, nn*, and *m* are integers; *nn* ∈ {01, 02, 03, 04, 05, 06, 07, 10, 11, 13, 15, 17, 18, 19} identifies the 15 individuals; and *m* ∈ {3, 4, 5, 6, 7} identifies the five time points. Files were loaded into R as a data frame using the `read.csv` command. Data frames were bound into a single data frame using the `cbind` command in R. For the methylation profiles, the GSE161020_series_matrix.txt.gz dataset, available in the GEO Series Matrix File(s) section, was used as is. The file was loaded into R as a data frame using the `read.csv` command. The first column of the data frame is an identifier in the form of cg*yyyyyyyy*, where *yyyyyyyy* is an integer. Since the methylation profile was measured with microarray technology, the identifier can be annotated with the reference to the microarray annotation file, GPL6480-9577.txt.gz, which is available under GEO ID GPL6480. Since the other 75 column names are in the form of GSM*XXXXXXX*, the columns were reordered with reference to the columns of the data frame generated from the gene expression profiles as described above. The proteome dataset was obtained from ProteomeX-change [11] using ID PXD020474. Two files (GR01,04,09,10,11,13,15,17,18,19.txt and GR02,03,05,06,07.txt) were downloaded and loaded as data frames into R using the read.csv command. The fourth column of the data frame includes the protein IDs, which were used as identifiers for subsequent analysis. These two data frames were merged with the row names of the union of the protein identifier. Missing observations were filled with zeros. Since the first and second rows have the format “GR*nn*” and “Visit *m*,” respectively, the columns can also be reordered with reference to the column names of the data frame generated by gene expression profiles as described above.

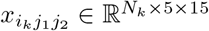 that is, the number of values of the number of *i*_*k*_ features of the *k*th feature type measured at the *j*_1_th time point for *j*_2_ individuals. These values are standardized as 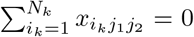 and 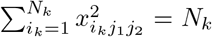 (*k* = 1 : methylation), 35829 (*k* = 2 : gene expression), 1588 (*k* = 3 : WBC proteome and *k* = 4 : plasma proteome). Figure 1 schematically illustrates the analysis method for the HBV vaccination datasets.

**Figure 1.**
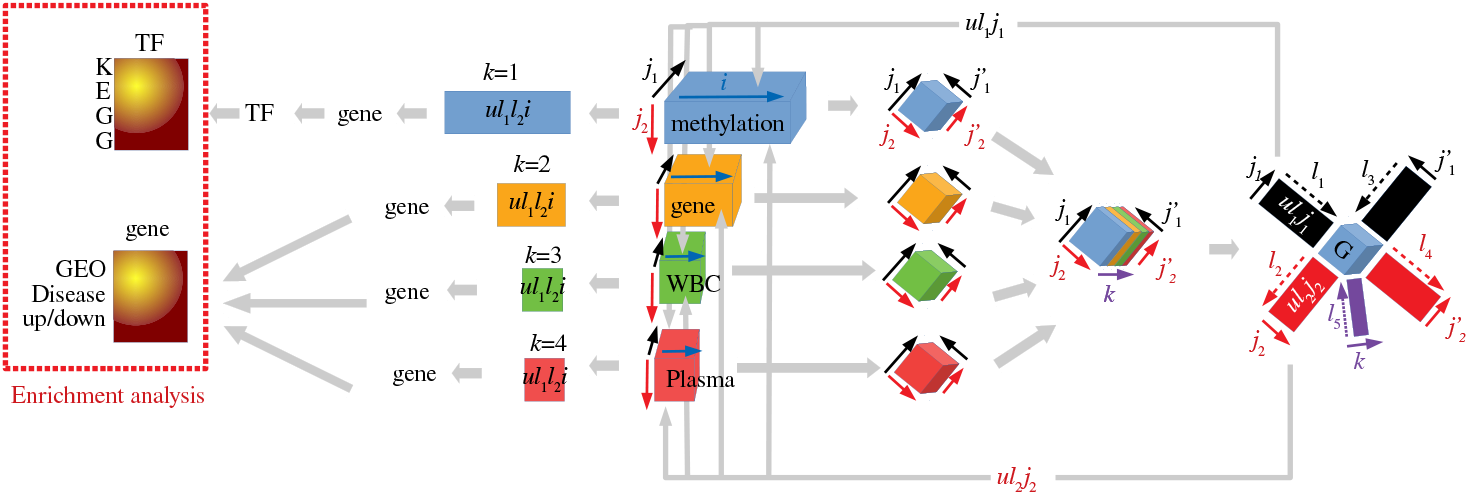
Schematic representation of hepatitis B virus (HBV) vaccination data analysis. Analysis starts from the center, moves to the right, comes back to the center, and then moves to the left. The cyan rectangle annotated as “methylation” is 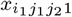, the yellow rectangle annotated as “gene” is 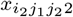, the green rectangle annotated as “WBC” is 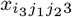, and the magenta rectangle annotated as “Plasma” is 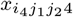. The four tilted cubes to the right of these four rectangles are 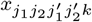, whose correspondence with 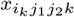 is indicated by the same color. The tilted cubes colored by layers to the right of the four tilted cubes represent the bundle of 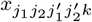. The right-most figure with a blue cube annotated as “G” at the center corresponds to TD shown in eq. (16). The four colored rectangles to the left of the four colored and annotated rectangles represent the singular-value vectors computed by eq. (17). Genes are selected from these singular-value vectors using *P*-values computed by eq. (18). For methylation, transcription factors (TFs) are further selected by Enricher using the selected genes (Table 3). The selected genes and TFs are then uploaded to Enrichr to validate the biological reliability (the left-most figure with color gradation).

A linear kernel was employed as follows:

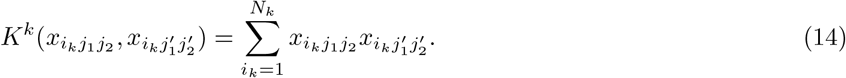

Then, a tensor was added:

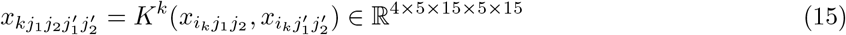

where 4 stands for four omics data, 5 stands for five time points and 15 stands for fifteen individuals. Applying HOSVD to 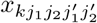 results in

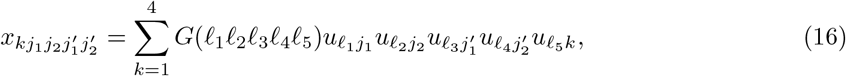

where 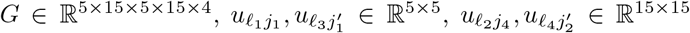, and 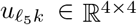. Since 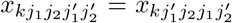 and 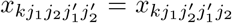 because of symmetry, 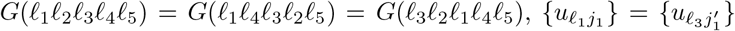, and 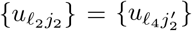.

The singular-value vectors of interest were as follows:

- 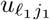 and 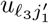 should be significantly dependent on time points corresponding to *j*_1_ and 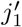.
- 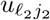 and 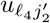 should be independent of individuals corresponding to *j*_2_ and 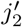.
- 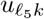 should be common between distinct omics measurements.

As a result, *ℓ*_1_ = *ℓ*_3_ = 2, *ℓ*_2_ = *ℓ*_4_ = 1, *ℓ*_5_ = 1 satisfies the required conditions (Fig. 2). Then,

**Figure 2.**
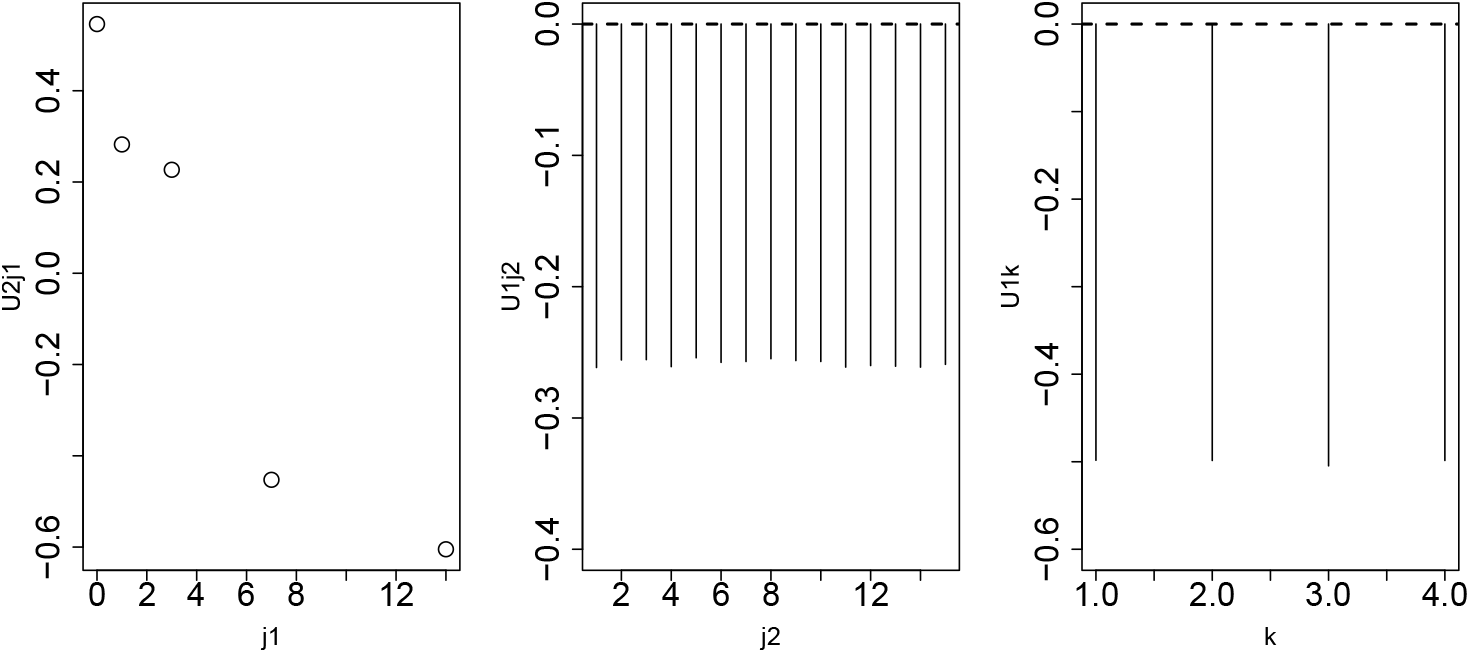
Left: 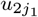, middle: 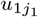, right *u*_1*k*_ when HOSVD is applied to a linear kernel computed using hepatitis B virus (HBV) vaccine data. The Pearson correlation coefficient between *j*_1_ and 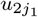 is *−*0.94 (*P* = 0.02).

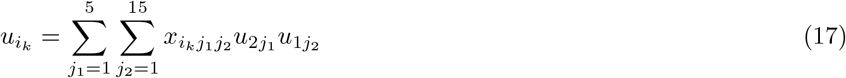

is computed and

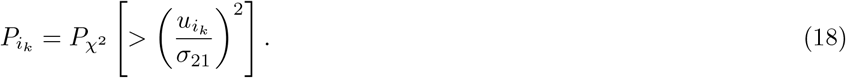

The computed 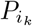 values were corrected using the BH criterion and *i*_*k*_s associated with either 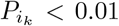 (for gene expression and methylation) or with 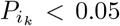 (for the two proteomes) were successfully selected (the full lists of selected features are available in Supplementary Data S1—S4).

### Kidney cancer multi-omics datasets

Full description of the compilation of the kidney cancer multi-omics datasets is available in the related study [12]. In brief, there were two sets of multi-omics kidney cancer data, each of which was composed of messenger RNA (mRNA) and microRNA (miRNA) expression profiles. The first dataset was obtained from The Cancer Genome Atlas (TCGA) and included 253 kidney tumors and 71 normal kidneys. The second dataset was obtained from GEO (GSE16441), and included 17 patients and 17 healthy controls. The method by which these two datasets were preprocessed is described in [12]. The dataset was formatted as 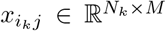, which represents The *i*_*k*_ expression levels of *j* subjects (*k* = 1 for mRNA and *k* = 2 for miRNA).

A linear kernel was employed as

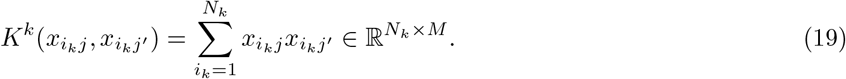

For the first dataset (i.e., TCGA data), *M* = 324, whereas for the second dataset (i.e., GEO data), *M* = 37. Then, a tensor was added

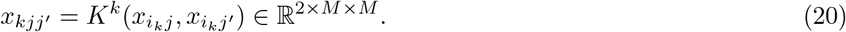

Applying HOSVD to *x*_*kjj’*_ results in

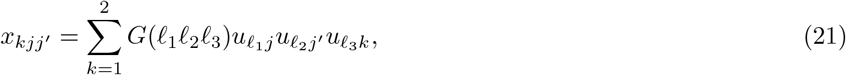

where 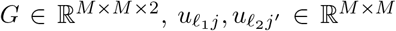, and 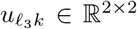. Since *x*_*kjj*′_ = *x*_*kj*′*j*_ because of symmetry, 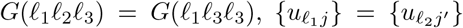. Then, singular-value vectors were identified such that 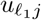 and 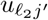 were significantly distinct between healthy controls and patients. As a result, *ℓ*_1_ = *ℓ*_2_ = 2 satisfies the required conditions (Fig. 3). Then,

**Figure 3.**
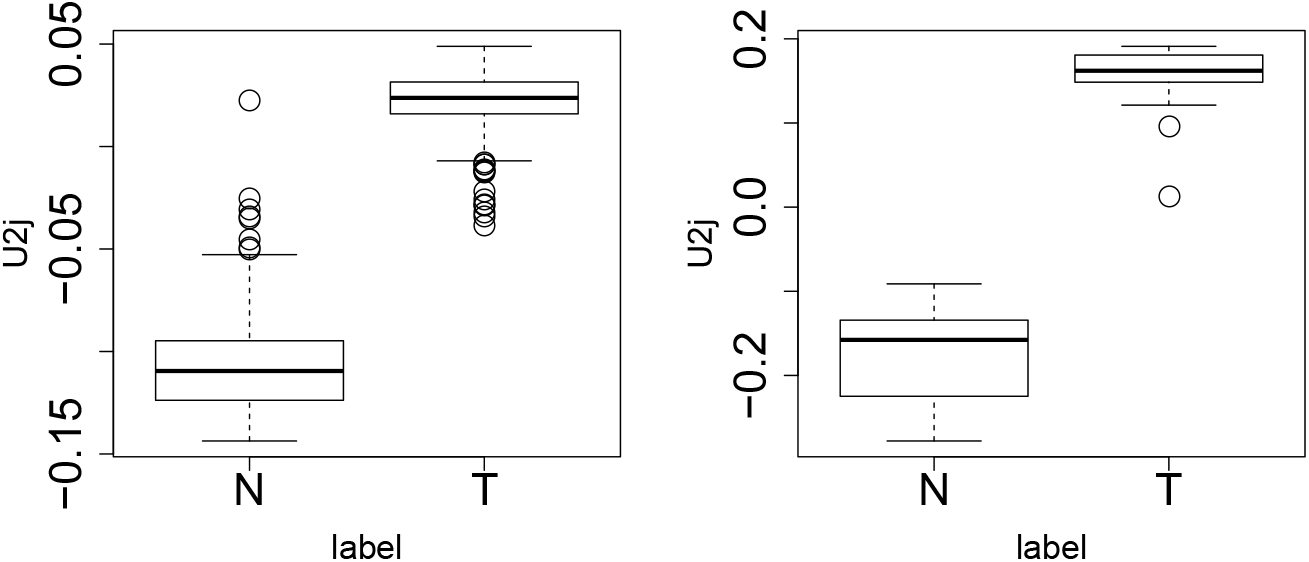
Boxplot of *u*_2*j*_ when higher-order singular-value decomposition (HOSVD) is applied to a linear kernel computed using kidney cancer data. Left: TCGA, *P* = 8.49*×* 10^*−*47^, right: GEO, *P* = 4.07 *×*10^*−*17^. *P*-values are based on the *t* test applied to *u*_2*j*_. T: tumors, N: normal kidney samples.

**Figure 4.**
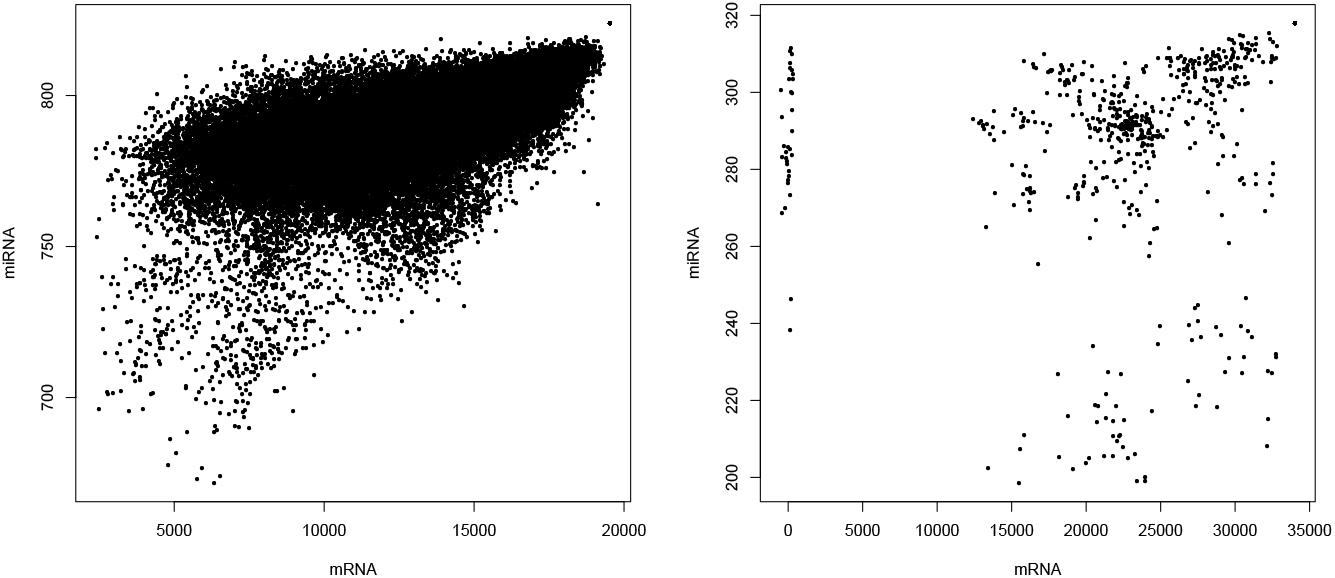
Scatter plot of kernels *K*^*k*^ between messenger RNA (mRNA) and microRNA (miRNA). Upper: The Cancer Genome Atlas (TCGA) (Pearson correlation coefficient = 0.627, *P* = 0.00), Lower: Gene Expression Omnibus (GEO)(Pearson correlation coefficient = 0.349, *P* = 2.32 *×* 10^*−*34^).

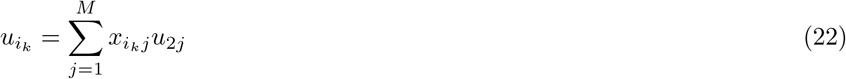

is computed and

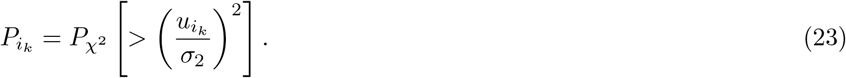

The computed 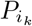 values were corrected using the BH criterion and 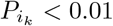 associated with *[ineq* were successfully selected.

## RESULTS

### Synthetic dataset

Table 1 shows the confusion matrix obtained by the proposed KTD-based unsupervised FE. Among the 10 features associated with *a*_*i*_, approximately seven features were correctly selected, whereas false positives were almost zero. Thus, KTD-based unsupervised FE successfully selected features correlated with *a*_*j*_.

Table 2 shows the confusion matrix obtained by linear regression-based FE. Essentially, no features correlated with *a*_*j*_ were selected. Thus, regression analysis did not select any features correlated with *a*_*j*_.

**Table 2.** Confusion matrix when linear regression, lasso and rf were applied to the synthetic dataset (*N* = 1000, *N*_1_ = 10, *M* = 10, *K* = 3). For cases when rf was employed, the results when the top most 2*N*_1_ features with larger absolute importance were selected have also been shown in parentheses.

|  | linear regression |  | lasso |  | rf |  |
| --- | --- | --- | --- | --- | --- | --- |
| | adjusted<br>$P_i \leq 0.01$ | adjusted<br>$P_i > 0.01$ | selected | not selected | selected | not selected |
| $k = 1$ | | | | | | |
| $i \leq 2N_1$ | 0.07 | 19.93 | 4.62 | 15.383 | 17.55 (5.82) | 2.45 (14.18) |
| $i > 2N_1$ | 0.03 | 979.97 | 2.12 | 977.88 | 495.43 (14.18) | 484.57 (965.82) |
| $k = 2$ | | | | | | |
| $i \leq N_1, 2N_1 < i \leq 3N_1$ | 0.07 | 19.93 | 4.70 | 15.30 | 17.69 (5.67) | 2.31 (14.33) |
| Other than above | 0.01 | 979.99 | 2.27 | 977.73 | 494.70 (14.33) | 485.30 (965.67) |
| $k = 3$ | | | | | | |
| $i \leq N_1, 3N_1 < i \leq 4N_1$ | 0.09 | 19.91 | 4.55 | 15.45 | 17.71 (5.46) | 2.29 (14.54) |
| Other than above | 0.01 | 979.99 | 2.12 | 977.78 | 496.68 (14.54) | 483.32 (965.46) |

**Table 3.** Transcription factors (TFs) enriched in the “ChEA 2016” Enrichr category (adjusted *P*-values *<* 0.05) when 1335 genes associated with 2077 methylation probes selected by KTD-based unsupervised FE were considered (the full list is available in Supplementary Data S1).

|  |  |
| --- | --- |
| TFs | ZNF217, TCF4, STAT3, SMARCD1, WT1, FOXA2, PAX3-FKHR, SMAD4, SMAD3, SOX9, TFAP2C, YAP1, AR, SOX2, CTNNB1, VDR, PIAS1, TEAD4, MITF, HNF4A, SUZ12 |

These results demonstrated that an apparently simple and easy problem became difficult when it is a *large p small n* problem, whereas KTD-based unsupervised FE was able to handle this problem to some extent. These advantages have also been observed in PCA- and TD-based unsupervised FE [4].

Although the methods that did not attribute *P*-values to features were not of interest, since the capability of attributing *P*-values to features is a great advantage of KTD-based unsupervised FE, as emphasized in the Background, lasso was employed as another method for comparison. Although lasso regression does not attribute *P*-values to features, the model was fitted to feature selection in a *large p small n* problem to demonstrate the difficulty of feature selection in the synthetic dataset. Table 2 shows the confusion matrix, which is clearly inferior to that shown in Table 1. Although the KTD-based unsupervised FE approach correctly selected at least 7 out of 10 features (70 %), which was correlated with *a*_*j*_ with essentially no false positives, lasso selected at most 5 out of 20 features (only 25 %), which was correlated with *a*_*j*_ with two false positives (approximately half of the true positives). In addition to lasso, we also tested rf as an alternative method that cannot attribute *P*-values (Table 2). First, we have selected features with nonzero importance; although most of (c.a. 17) features among the 20 features coincident with *a*_*j*_ are selected, almost half of features not coincident with *a*_*j*_ are also wrongly selected. Thus, rf is clearly inferior to TD-based unsupervised FE. Although one might wonder if the top 20 features with a larger absolute importance are selected, only five out of 20 features are selected (Table 2). Thus, rf is still inferior to TD-based unsupervised FE even if limited and top most important features are selected. This suggested that even if methods that could not attribute *P*-values to features can be considered, they would not outperform the KTD-based unsupervised FE method. Thus, subsequently, we only focused only methods that attributed *P*-values to features.

### HBV vaccine dataset

To validate genes selected by KTD-based unsupervised FE, the selected genes were uploaded to Enrcihr [13]. Initially, 1335 genes associated with 2077 methylation probes selected by KTD-based unsupervised FE were uploaded. Many transcription factors (TFs) were significantly predicted to target these 1335 genes (Table 3). These 21 TFs were then uploaded to Enrichr again; Table S1 shows the top 10 Kyoto Encyclopedia of Genes and Genomes (KEGG) pathways in the “KEGG 2019 HUMAN” Enrichr category (the full list is available in Supplementary Data S1). It was clear that these data included many biologically reasonable KEGG pathways (for details, see the Discussion section below).

Eight genes associated with 11 probes were identified as differentially expressed genes (DEGs) when gene expression profiles were considered (Table 4), which were uploaded to Enrichr. Conversion of probe names to gene symbols was performed by the ID converter tool of DAVID [14]. Table S2 shows the top 10 GEO profiles enriched in the Enrichr “Disease Perturbations from GEO up/down” category (the full list is available in Supplementary Data S2). Although many neurodegenerative profiles not apparently related to the HBV vaccine are listed, the rationalization for their inclusion is provided in the Discussion section.

**Table 4.** Eight genes associated with 11 probes identified as differentially expressed genes (DEGs) when gene expression profiles were considered. Proteins identified as differentially expressed genes (DEGs) when gene expression profiles in the proteome were considered.

|  |  |
| --- | --- |
| Gene symbols | S100A9, CD74, hba1, ACTB, HBB, HBA2, MALAT1, COX1 |
| WBC | HIST1H2BJ, HIST2H2BF, HIST1H2BG, HIST1H2BB, HIST1H2BD, ACTG1, HIST1H2BL, HIST1H2BN, PFN1, HIST1H2BK, HIST3H2BB, ACTB, HBB, HBA2, HIST1H2BA, HIST1H2BI, HIST1H2BC, HIST1H2BO, HIST2H2BE, HIST1H2BM, HBA1, HIST1H2BF, HIST1H2BE, HIST1H2BH |
| Plasma | FGA, HP, GSN, ALB, FGG, IGLL5, APOA1, SERPINA1, ORM1, TF, GC, CP, C4A, CSF3R, A2M, HPX, HRG, A1BG, CFH, APOB, C3, CLEC14A |

Finally, two set of proteins identified as DEGs in WBC and plasma profiles (Table 4) were uploaded to Enrichr. Tables S3 and S4 show the top 10 enriched GEO profiles identified in the Enrichr “Disease Perturbations from GEO up/down” category when proteins for the WBC and plasma sections in Table 5 were uploaded to Enrichr (the full list is available in Supplementary Data S3 and S4). Other than the enriched reasonable hepatitis-related GEO profiles, some neurodegenerative disease-related GEO profiles were also enriched, as shown in Table S3, which are further explored in the Discussion section.

**Table 5.** Proteins identified as differentially expressed genes (DEGs) when gene expression profiles in the proteome were considered.

|  |  |
| --- | --- |
| WBC | HIST1H2BJ, HIST2H2BF, HIST1H2BG, HIST1H2BB, HIST1H2BD, ACTG1, HIST1H2BL, HIST1H2BN, PFN1, HIST1H2BK, HIST3H2BB, ACTB, HBB, HBA2, HIST1H2BA, HIST1H2BI, HIST1H2BC, HIST1H2BO, HIST2H2BE, HIST1H2BM, HBA1, HIST1H2BF, HIST1H2BE, HIST1H2BH |
| Plasma | FGA, HP, GSN, ALB, FGG, IGLL5, APOA1, SERPINA1, ORM1, TF, GC, CP, C4A, CSF3R, A2M, HPX, HRG, A1BG, CFH, APOB, C3, CLEC14A |

Since there are many enriched biological processes and pathways listed in Tables S1–S4, the genes and proteins selected in this section may not be artifacts, but likely have a true biological basis.

### Kidney cancer

hsa-mir-200c and hsa-mir-141 were selected from TCGA, and hsa-miR-141, hsamiR-210, and hsa-miR-200c were selected from GEO. Thus, these miRNAs are highly coincident with each other, even more so than reported in previous works [12, 5] where TD-as well as KTD-based unsupervised FE was applied to TCGA and GEO datasets. For mRNA, there were five common genes selected between the TCGA and GEO datasets (Table 6, *P* = 6.7 × 10^*−*5^, odds ratio: 13.13).

**Table 6.** Confusion matrix of selected mRNAs between The Cancer Genome Atlas (TCGA) and Gene Expression Omnibus (GEO) datasets

|  |  | GEO |  |
| --- | --- | --- | --- |
| | | $P > 0.01$ | $P < 0.01$ |
| TCGA | $P > 0.01$ | 17269 | 101 |
| | $P < 0.01$ | 65 | 5 |
| | | $P = 6.7 \times 10^{-5}$, Odds ratio: 13.13 | |

## DISCUSSION

There are several advantages in the proposed implementation of KTD-based unsupervised FE compared with previously proposed versions of KTD-based unsupervised FE [5], as well as TD-based unsupervised FE [4] in the context of application to the integration of multi-omics datasets. For example,

1. The present implementation can reduce the required computational memory. As a result, required computational time can be reduced as well.
2. The present implementation can integrate more than two omics data in a straight manner.

Although the point was achieved in KTD-based unsupervised FE [5], too, the primary advantage of the proposed method is to achieve the above two simultaneously, as discussed below. As for point 1, when the original implementation of TD-based unsupervised FE [4] is applied to the integration of multi-omics data, (e.g., *x*_*ij*_ ∈ ℝ ^*N*×*M*^ and *x*_*hj*_ ∈ ℝ ^*H*×*M*^, which corresponds to the *i*th or *h*th omics data of the *j*th sample), HOSVD is applied to

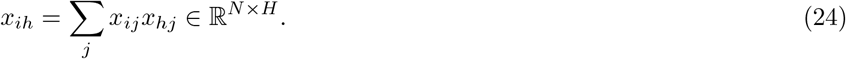

Since *H, N* » *M*, this was not an effective implementation. When KTD-based unsupervised FE [5] is applied, HOSVD is applied to

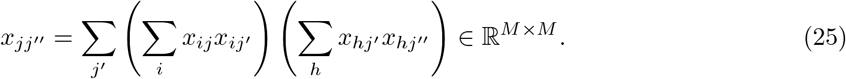

This drastically reduced the required memory and central processing unit (CPU) time. In the present implementation. we can keep this advantage as well, since the size of tensors used in this study is independent of the number of features, *N*. As for point 2, nevertheless, it was unclear how more than two omics datasets could be integrated. In the implementation introduced in this paper,

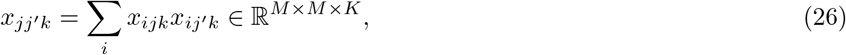

which corresponds to the *i*th measurement of the *k*th omics data of the *j*th sample, where *K* is the total number of omics datasets. Since HOSVD is applied to *x*_*jj*′*k*_, any number of multi-omics datasets may be handled. This slight modification drastically increased the ability of KTD-based unsupervised FE to handle multi-omics datasets. Thus, it is obvious that the present implementation has at least one advantage over the past KTD implementations.

Kernels *K*^*k*^ were highly correlated between mRNA (*k* = 1) and miRNA (*k* = 2) for the kidney cancer data [15] (Fig.4). Kernels *K*^*k*^ for gene expression profiles and the proteome were also highly correlated when HBV vaccine experiments were considered (Table 7). Thus, it was obvious that the current formalism was very effective in identifying the coincidence between individual omics (in this case, mRNA, miRNA, and the proteome).

**Table 7.**
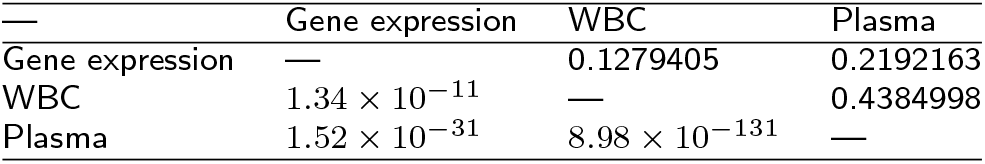
Correlation between hepatitis B virus (HBV) vaccine experiment kernels. Upper triangle: Pearson correlation, lower triangle: *P*-values.

Although the list of enriched pathways in Table S1 did not seem to be related to the HBV vaccine directly, there were indirect reasonable relationships. For example, the Hippo signaling pathway has recently been identified to be related to the immune system [16]. “Hepatitis B” was also significantly enriched (adjusted *P*-value of 7.79× 10^*−*3^), but it did not rank within the top 10 KEGG pathways in Table S1. In addition, since patients with diabetes have a higher risk of HBV infection [17], it is reasonable that the two KEGG pathways “AGE-RAGE signaling pathway in diabetic complications” and “Maturity onset of diabetes in the young” were enriched. Although multiple cancer types were also enriched in these data, as shown in Table S1, many cancer types other than liver cancer are known to be related to the risk of HBV infection [18].

Although genes identified as DEGs in relation to HBV vaccination were also enriched in various neurodegenerative diseases other than hepatitis (Tables S2 and S3), this is a reasonable finding because viral hepatitis was reported to be related to Parkinson’s disease [19]. There are also known associations between hepatic functions and plasma amyloid-*β* levels [20]; cirrhosis patients with HBV infection have higher plasma A*β*40 and A*β*42 levels than patients with HBV-negative cirrhosis. More directly, Ji et al.[21] reported that the hepatitis B core VLP-based mis-disordered tau vaccine alleviated cognitive deficits and neuropathology progression in a Tau.P301S mouse model of Alzheimer’s disease. Thus, enrichment of neurodegenerative disease-related genes among the identified DEGs does not appear to be an artifact, but rather provides possible supportive evidence that KTD-based unsupervised FE detected side effects caused by vaccinations.

Other conventional univariate tools such as limma [22] and sam [23] cannot be used for these tasks since they are designed to handle categorical classes and thus cannot be applied to HBV vaccination data, which are only associated with time points and are not categorical. Although regression analysis was attempted for the synthetic dataset, there were no features correlated with dates. Thus, there were no univariate feature selection methods applicable to the HBV vaccination data that could identify features correlated with date. For the kidney cancer datasets, it has been extensively demonstrated that these conventional univariate tools such as limma [22] and sam [23] cannot compete with the TD-based unsupervised FE approach [12, 5]. Thus, no univariate feature selection method was identified that was superior to TD-based unsupervised FE when applied to kidney cancer datasets.

The performance of the proposed method has not been compared to other existing multi-omics–oriented methods [1, 2] because no suitable methods were identified for suitable comparison. First, most of the recently proposed cutting-edge methods adapted to multi-omics analysis are specific to a high-throughput sequencing (HTS) architecture. For example, MKpLMM [24] requires genomic coordinates, which are not available for the datasets analyzed in this study. Similarly, csaw [25] requires a bed file, which is also not available for the present dataset. Since the purpose of the present study was to propose a more flexible method that is not specific to the HTS architecture and the datasets employed in this study were not obtained using HTS, these were considered unsuitable methods for comparison with the proposed implementation of KTD-based unsupervised FE. Second, other methods that are not specific to HTS lack the statistical validation of feature selection (i.e., no ability to attribute *P*-values to features). For example, although MOFA [26] is not specific to HTS, it does not have the ability to select features; thus, we were not able to compare its performance with that of our proposed method. Although DIABLO [27] is also not specific to the HTS architecture and has feature selection ability, there is no functionality to attribute *P*-values to individual features; thus, features cannot be selected based on statistical significance, and therefore, DIABLO was considered to be outside of the scope of this study. FSMKL [28], which is also not specific to the HTS architecture does have the ability to add statistical scores to individual features; however, selecting features based on statistical scores is not effective. In this sense, to our knowledge, there are no other multi-omics–oriented feature selection methods that satisfy the following requirements:

- Not specific to the HTS architecture
- Attributes *P*-values to individual features to evaluate statistical significance
- Can handle more than or equal to three kinds of omics data simultaneously
- Applicable to severe *large p small n* (typically, *p/n* ∼ 10^2^ or more) problems For example, for the *large p small n* problem, although Subramanian et al.[2] summarized existing machine learning methods for multi-omics analysis, typically they are applied to studies including up to 10^2^ samples; thus, they cannot be regarded as a severe *large p small n* problem. As a result, the performance of the KTD-based unsupervised FE was not directly compared to other existing methods that satisfy all of the above conditions.

HBV vaccination data were selected to demonstrate the superior power of advanced KTD-based unsupervised FE because of the difficulty of the problem with this dataset. Since vaccination must be given to healthy people, side effects must be minimized [29]; in fact, since vaccination is essentially an infection with a weaker pathogen, its effect is inevitably weak. As expected, a very limited number of features (genes and proteins) were selected using the advanced KTD-based unsupervised FE proposed in this article, whereas conventional linear regression analysis did not attribute significant *P*-values to any features. This suggests that the proposed advanced KTD-based unsupervised FE method has superior ability to select features when applied to even particularly difficult multi-omics datasets.

One might wonder why we did not employ more advanced feature selection methods other than lasso or rf. Generally, other methods are not fitted to the present situation, i.e., the *large p small n* problem. Since it is impossible to demonstrate the difficulty of using all the other methods, we consider two methods, class-specific mutual information variation for feature selection [30] and multilabel feature selection with constrained latent structure shared term [31] in order to demonstrate why other advanced methods are not fitted to the *large p small n* problem. When *k* features were aimed to be selected, although the complexity of class-specific mutual information variation for feature selection was supposed to be *kMN*, it excluded the computational time needed for the computation of mutual information among *N* features, which is as large as *N* ^2^. It is not fitted to the *large p small n* problem associated with a large number of features, *N*. For example, in the synthetic example, we tried to compute mututal information among *N* = 100 features for only one ensemble; it took 100 seconds. Since we employed *N* = 1000, it would take 100 × (1000*/*100)^2^ = 10^4^ seconds for only one ensemble. We employed 100 ensembles; thus, in total, the required computational time for 100 ensembles would be as long as 10^4^ × 100 = one million seconds, which is unrealistic, since other methods, such as linear regression, lasso and rf, require less than 10 minutes (= 600 seconds) for computation with 100 ensembles. This means that class-specific mutual information variation for feature selection is not reasonable to be applied to the present synthetic example. As for multilabel feature selection with constrained latent structure shared term, it is not fitted to the *large p small n* problem as well, since it can select at most *M* features. Multilabel feature selection with constrained latent structure shared term decomposes the matrix *N* × *M* into a product of two small matricies, *N* × *k* and *k* × *M*, when *k* features are selected. Nevertheless, in the *large p small n* problem, since *M* « *N, k < M < N*. Thus, it can select at most *M* features. On the other hand, multilabel feature selection with constrained latent structure shared term was applied to the case where *N < M* [31]. In our synthetic example, the number of features to be selected, 2*N*_1_, is larger than *M*. Thus, multilabel feature selection with constrained latent structure shared term cannot be used for the present synthetic example. Although these are only two examples, most of the popular feature selection methods are not suitable for the *large p small n* problem, as shown for these two methods.

## CONCLUSION

In this paper, an advanced KTD-based unsupervised FE method was introduced, which was modified to be applied to feature selection in multi-omics data analysis that is often very difficult, mainly based on the *large p small n* problem. The proposed method was successfully applied to a synthetic dataset, as well as to two real datasets, and attributed significant *P*-values to features with reduced CPU time and memory, even when applied to integrated analysis of more than two multi-omics datasets. Although the modification from the previously proposed KTD-based unsupervised FE was not significant, this slight modification was successful when applied to feature selection of multi-omics data analysis, which often poses a challenge in the case of a *large p small n* problem.

## Declarations

### Ethics approval and consent to participate

Not applicable.

### Consent for publication

Not applicable.

### Availability of data and materials

TCGA data set can be downloaded from http://firebrowse.org/ (No accession number was assigned). GEO data set can be downloaded using GEO ID: GSE16441.

### Author’s contributions

YHT planned the research, performed analyses. YHT and TT have evaluated the results, discussions, outcomes and wrote and reviewed the manuscript.

## Acknowledgements

Not applicable.

## Additional Files

Additional file 1 — Supplementary Tables Tables S1 to S4

Additional file 2 — Supplementary Data S1

Data_S1.xlsx, list of selected features and enrichment analysis for methylation.

Additional file 3 — Supplementary Data S2

Data_S2.xlsx, list of selected features and enrichment analysis for gene expression,

Additional file 4 — Supplementary Data S3

Data_S3.xlsx, list of selected features and enrichment analysis for proteome of WBC.

Additional file 5 — Supplementary Data S4

Data_S4.xlsx, list of selected features and enrichment analysis for proteome of Plasma.

## References

1. Reel, P.S., et al.: Using machine learning approaches for multi-omics data analysis: A review. Biotechnology Advances 49, 107739 (2021)

2. Subramanian, I., et al.: Multi-omics data integration, interpretation, and its application. Bioinformatics and Biology Insights 14, 1177932219899051 (2020)

3. Huynh, P.-H., et al.: Improvements in the large p, small n classification issue. SN Computer Science 1(4) (2020)

4. Taguchi, Y.-h.: Unsupervised Feature Extraction Applied to Bioinformatics. Springer, ??? (2020)

5. Taguchi, Y.-H., Turki, T.: Mathematical formulation and application of kernel tensor decomposition based unsupervised feature extraction. Knowledge-Based Systems 217, 106834 (2021)

6. Roy, S.S., Taguchi, Y.-H.: Identification of genes associated with altered gene expression and m6a profiles during hypoxia using tensor decomposition based unsupervised feature extraction. Scientific Reports 11(1), 8909 (2021)

7. Taguchi, Y.-H.: Tensor decomposition-based and principal-component-analysis-based unsupervised feature extraction applied to the gene expression and methylation profiles in the brains of social insects with multiple castes. BMC Bioinformatics 19(S4) (2018). doi:10.1186/s12859-018-2068-7

8. R Core Team: R: A Language and Environment for Statistical Computing. R Foundation for Statistical Computing, Vienna, Austria (2020). R Foundation for Statistical Computing. https://www.R-project.org/

9. Tibshirani, R.: Regression shrinkage and selection via the lasso. Journal of the Royal Statistical Society. Series B (Methodological) 58(1), 267–288 (1996)

10. Liaw, A., Wiener, M.: Classification and regression by randomforest. R News 2(3), 18–22 (2002)

11. Deutsch, E.W., et al.: The ProteomeXchange consortium in 2017: supporting the cultural change in proteomics public data deposition. Nucleic Acids Research 45(D1), 1100–1106 (2016)

12. Ng, K.-L., Taguchi, Y.-H.: Identification of miRNA signatures for kidney renal clear cell carcinoma using the tensor-decomposition method. Scientific Reports 10(1) (2020)

13. Kuleshov, M.V., et al.: Enrichr: a comprehensive gene set enrichment analysis web server 2016 update. Nucleic Acids Research 44(W1), 90–97 (2016)

14. Huang, D.W., et al.: Systematic and integrative analysis of large gene lists using DAVID bioinformatics resources. Nature Protocols 4(1), 44–57 (2008)

15. Li, X., et al.: Integrated analysis of MicroRNA (miRNA) and mRNA profiles reveals reduced correlation between MicroRNA and target gene in cancer. BioMed Research International 2018, 1–15 (2018)

16. Hong, L., et al.: Role of hippo signaling in regulating immunity. Cellular & Molecular Immunology 15(12), 1003–1009 (2018)

17. Khalili, M., et al.: Diabetes and prediabetes in patients with hepatitis b residing in north america. Hepatology 62(5), 1364–1374 (2015)

18. Song, C., OTHERS: Associations Between Hepatitis B Virus Infection and Risk of All Cancer Types. JAMA Network Open 2(6), 195718–195718 (2019)

19. Pakpoor, J., et al.: Viral hepatitis and parkinson disease. Neurology 88(17), 1630–1633 (2017)

20. Wang, Y.-R., et al.: Associations between hepatic functions and plasma amyloid-beta levels—implications for the capacity of liver in peripheral amyloid-beta clearance. Molecular Neurobiology 54(3), 2338–2344 (2016)

21. Ji, M., et al.: Hepatitis B core VLP-based mis-disordered tau vaccine elicits strong immune response and alleviates cognitive deficits and neuropathology progression in tau.p301s mouse model of alzheimer’s disease and frontotemporal dementia. Alzheimer’s Research & Therapy 10(1) (2018)

22. Ritchie, M.E., et al.: limma powers differential expression analyses for RNA-sequencing and microarray studies. Nucleic Acids Research 43(7), 47 (2015)

23. Tusher, V.G., et al.: Significance analysis of microarrays applied to the ionizing radiation response. Proceedings of the National Academy of Sciences 98(9), 5116–5121 (2001)

24. Li, J., et al.: Multi-kernel linear mixed model with adaptive lasso for prediction analysis on high-dimensional multi-omics data. Bioinformatics 36(6), 1785–1794 (2019)

25. Lun, A.T.L., Smyth, G.K.: csaw: a Bioconductor package for differential binding analysis of ChIP-seq data using sliding windows. Nucleic Acids Research 44(5), 45–45 (2015)

26. Argelaguet, R., et al.: Multi-omics factor analysis-a framework for unsupervised integration of multi-omics data sets. Molecular Systems Biology 14(6), 8124 (2018)

27. Singh, A., et al.: DIABLO: an integrative approach for identifying key molecular drivers from multi-omics assays. Bioinformatics 35(17), 3055–3062 (2019)

28. Seoane, J.A., et al.: A pathway-based data integration framework for prediction of disease progression. Bioinformatics 30(6), 838–845 (2013)

29. Jacobson, R.M., et al.: Making vaccines more acceptable - methods to prevent and minimize pain and other common adverse events associated with vaccines. Vaccine 19(17), 2418–2427 (2001)

30. Gao, W., Hu, L., Zhang, P.: Class-specific mutual information variation for feature selection. Pattern Recognition 79, 328–339 (2018). doi:10.1016/j.patcog.2018.02.020

31. Gao, W., Li, Y., Hu, L.: Multilabel feature selection with constrained latent structure shared term. IEEE Transactions on Neural Networks and Learning Systems, 1–10 (2021). doi:10.1109/TNNLS.2021.3105142

