## Supplementary Data S1--S4 for "Novel feature selection method via kernel tensor decomposition for improved multi-omics data analysis": Supplementary_Tables.pdf

Table S1. Top-ranked 10 Kyoto Encyclopedia of Genes and Genomes (KEGG) pathways enriched in the Enrichr "KEGG 2019 HUMAN" category when 21 transcription factors (TFs) in Table 4 are uploaded. (The full list is available in Supplementary Data S1.)

| Term | Overlap | P-value | Adjusted P-value |
|---|---|---|---|
| Signaling pathways regulating pluripotency of stem cells | 6/139 | $4.21 \times 10^{-8}$ | $3.16 \times 10^{-6}$ |
| Hippo signaling pathway | 6/160 | $9.73 \times 10^{-8}$ | $3.65 \times 10^{-6}$ |
| Pathways in cancer | 8/530 | $6.06 \times 10^{-7}$ | $1.52 \times 10^{-5}$ |
| Th17 cell differentiation | 4/107 | $1.66 \times 10^{-5}$ | $3.11 \times 10^{-4}$ |
| Hepatocellular carcinoma | 4/168 | $9.68 \times 10^{-5}$ | $1.45 \times 10^{-3}$ |
| Adherens junction | 3/72 | $1.53 \times 10^{-4}$ | $1.85 \times 10^{-3}$ |
| Pancreatic cancer | 3/75 | $1.73 \times 10^{-4}$ | $1.85 \times 10^{-3}$ |
| Colorectal cancer | 3/86 | $2.59 \times 10^{-4}$ | $2.43 \times 10^{-3}$ |
| AGE-RAGE signaling pathway in diabetic complications | 3/100 | $4.03 \times 10^{-4}$ | $3.36 \times 10^{-3}$ |
| Maturity onset diabetes of the young | 2/26 | $6.46 \times 10^{-4}$ | $4.84 \times 10^{-3}$ |

Table S2. Top-ranked 10 Gene Expression Omnibus (GEO) profiles enriched in the Enrichr "Disease Perturbations from GEO up/down" category when 8 genes in Table 5 are uploaded (the full list is available in Supplementary Data S2).

| Disease Perturbations from GEO down | | | |
|---|---|---|---|
| Term | Overlap | P-value | Adjusted P-value |
| Parkinson's disease DOID-14330 human GSE7621 sample 941 | 6/258 | $1.19 \times 10^{-10}$ | $5.31 \times 10^{-8}$ |
| Teratospermia UMLS CUI-C0919628 human GSE6872 sample 953 | 6/372 | $1.08 \times 10^{-9}$ | $2.41 \times 10^{-7}$ |
| autism spectrum disorder DOID-0060041 human GSE25507 sample 1029 | 5/297 | $3.77 \times 10^{-8}$ | $4.65 \times 10^{-6}$ |
| asthma DOID-2841 human GSE16032 sample 729 | 5/303 | $4.17 \times 10^{-8}$ | $4.65 \times 10^{-6}$ |
| familial hypercholesterolemia DOID-13810 human GSE6088 sample 908 | 5/317 | $5.22 \times 10^{-8}$ | $4.66 \times 10^{-6}$ |
| relapsing-remitting multiple sclerosis DOID-2378 human GSE16461 sample 879 | 5/359 | $9.71 \times 10^{-8}$ | $7.22 \times 10^{-6}$ |
| multiple sclerosis DOID-2377 human GSE16461 sample 584 | 5/452 | $3.05 \times 10^{-7}$ | $1.83 \times 10^{-5}$ |
| Fracture of femur C0015802 rat GSE1685 sample 240 | 4/168 | $3.28 \times 10^{-7}$ | $1.83 \times 10^{-5}$ |
| Malignant mesothelioma of pleura C0812413 human GSE2549 sample 118 | 4/203 | $6.99 \times 10^{-7}$ | $3.37 \times 10^{-5}$ |
| Parkinson's disease DOID-14330 human GSE19587 sample 740 | 4/207 | $7.55 \times 10^{-7}$ | $3.37 \times 10^{-5}$ |

| Disease Perturbations from GEO up | | | |
|---|---|---|---|
| Term | Overlap | P-value | Adjusted P-value |
| autism spectrum disorder DOID-0060041 human GSE25507 sample 1032 | 6/253 | $1.06 \times 10^{-10}$ | $5.72 \times 10^{-8}$ |
| facioscapulohumeral muscular dystrophy DOID-11727 human GSE15090 sample 541 | 5/162 | $1.80 \times 10^{-9}$ | $3.19 \times 10^{-7}$ |
| actinic keratosis DOID-8866 human GSE2503 sample 628 | 5/170 | $2.29 \times 10^{-9}$ | $3.19 \times 10^{-7}$ |
| Actinic keratosis C0022602 human GSE2503 sample 350 | 5/171 | $2.36 \times 10^{-9}$ | $3.19 \times 10^{-7}$ |
| melanoma in situ UMLS CUI-C0346040 human GSE4587 sample 980 | 5/196 | $4.70 \times 10^{-9}$ | $5.07 \times 10^{-7}$ |
| skin squamous cell carcinoma DOID-3151 human GSE2503 sample 627 | 5/227 | $9.82 \times 10^{-9}$ | $8.83 \times 10^{-7}$ |
| schizophrenia DOID-5419 human GSE21935 sample 855 | 5/250 | $1.59 \times 10^{-8}$ | $1.23 \times 10^{-6}$ |
| multiple sclerosis DOID-2377 human GSE26484 sample 742 | 5/271 | $2.38 \times 10^{-8}$ | $1.61 \times 10^{-6}$ |
| Schizophrenia C0036341 human GSE4036 sample 357 | 5/291 | $3.40 \times 10^{-8}$ | $1.87 \times 10^{-6}$ |
| Sjogren's syndrome DOID-12894 human GSE23117 sample 889 | 5/292 | $3.46 \times 10^{-8}$ | $1.87 \times 10^{-6}$ |

Table S3. Top-ranked 10 Gene Expression Omnibus (GEO) profiles enriched in the Enrichr "Disease Perturbations from GEO up/down" category when proteins listed in the WBC section of Table 6 are uploaded (the full list is available in Supplementary Data S3).

| Disease Perturbations from GEO down | | | |
|---|---|---|---|
| Term | Overlap | P-value | Adjusted P-value |
| ulcerative colitis DOID-8577 human GSE11223 sample 593 | 11/297 | $1.35 \times 10^{-14}$ | $4.99 \times 10^{-12}$ |
| autism spectrum disorder DOID-0060041 human GSE7329 sample 1034 | 11/308 | $2.02 \times 10^{-14}$ | $4.99 \times 10^{-12}$ |
| psoriasis DOID-8893 human GSE14905 sample 754 | 8/322 | $2.43 \times 10^{-9}$ | $4.01 \times 10^{-7}$ |
| asthma DOID-2841 human GSE16032 sample 729 | 7/303 | $4.74 \times 10^{-8}$ | $5.86 \times 10^{-6}$ |
| Non-syndromic cleft lip and palate DOID-9296 human GSE42589 sample 618 | 7/426 | $4.79 \times 10^{-7}$ | $4.73 \times 10^{-5}$ |
| autistic disorder DOID-12849 human GSE6575 sample 1042 | 5/143 | $6.64 \times 10^{-7}$ | $5.46 \times 10^{-5}$ |
| hepatitis C DOID-1883 human GSE20948 sample 600 | 6/307 | $1.33 \times 10^{-6}$ | $9.37 \times 10^{-5}$ |
| Hereditary gingival fibromatosis C0399440 human GSE4250 sample 177 | 6/321 | $1.72 \times 10^{-6}$ | $9.97 \times 10^{-5}$ |
| cystic fibrosis DOID-1485 mouse GSE33319 sample 1048 | 6/324 | $1.82 \times 10^{-6}$ | $9.97 \times 10^{-5}$ |
| Huntington's disease DOID-12858 human GSE1751 sample 795 | 5/213 | $4.71 \times 10^{-6}$ | $2.33 \times 10^{-4}$ |
| Disease Perturbations from GEO up | | | |
| Term | Overlap | P-value | Adjusted P-value |
| systemic juvenile idiopathic arthritis (sJIA) DOID-848 human GSE21521 sample 574 | 11/367 | $1.38 \times 10^{-13}$ | $3.91 \times 10^{-11}$ |
| acute myeloid leukemia DOID-9119 human GSE9476 sample 782 | 9/163 | $1.49 \times 10^{-13}$ | $3.91 \times 10^{-11}$ |
| Down syndrome DOID-14250 human GSE19681 sample 1066 | 10/280 | $4.06 \times 10^{-13}$ | $7.08 \times 10^{-11}$ |
| H1N1 DOID-0050211 human GSE27131 sample 514 | 11/427 | $7.19 \times 10^{-13}$ | $9.40 \times 10^{-11}$ |
| hepatocellular carcinoma DOID-684 human GSE58208 sample 735 | 10/455 | $4.96 \times 10^{-11}$ | $5.19 \times 10^{-9}$ |
| Lewy body dementia DOID-12217 human GSE49036 sample 1068 | 9/366 | $2.14 \times 10^{-10}$ | $1.87 \times 10^{-8}$ |
| Schistosomiases C0036323 mouse GSE19525 sample 439 | 9/485 | $2.55 \times 10^{-9}$ | $1.90 \times 10^{-7}$ |
| Huntington's disease DOID-12858 human GSE8762 sample 931 | 7/315 | $6.19 \times 10^{-8}$ | $3.76 \times 10^{-6}$ |
| polycystic ovary syndrome DOID-11612 human GSE10946 sample 825 | 7/320 | $6.89 \times 10^{-8}$ | $3.76 \times 10^{-6}$ |
| systemic juvenile idiopathic arthritis (sJIA) (enthesitis-related) DOID-848 human GSE21521 sample 575 | 7/322 | $7.19 \times 10^{-8}$ | $3.76 \times 10^{-6}$ |

Table S4. Top-ranked 10 Gene Expression Omnibus (GEO) profiles enriched in the Enrichr "Disease Perturbations from GEO up/down" category when proteins listed in the plasma section of Table 6 are uploaded (the full list is available in Supplementary Data S4).

| Disease Perturbations from GEO down | | | |
|---|---|---|---|
| Term | Overlap | P-value | Adjusted P-value |
| Nemaline Myopathy C0206157 mouse GSE3384 sample 160 | 12/324 | $1.49 \times 10^{-16}$ | $5.81 \times 10^{-14}$ |
| nemaline myopathy DOID-3191 mouse GSE3384 sample 976 | 12/450 | $7.68 \times 10^{-15}$ | $1.50 \times 10^{-12}$ |
| nemaline myopathy DOID-3191 mouse GSE3384 sample 975 | 10/239 | $2.80 \times 10^{-14}$ | $3.64 \times 10^{-12}$ |
| Hepatitis, Autoimmune C0241910 mouse GSE867 sample 230 | 8/227 | $6.79 \times 10^{-11}$ | $6.62 \times 10^{-9}$ |
| Carcinoma, Hepatocellular C0019204 human GSE6764 sample 407 | 8/303 | $6.73 \times 10^{-10}$ | $5.25 \times 10^{-8}$ |
| Gastrointestinal stromal tumor C0238198 human GSE2719 sample 270 | 6/224 | $1.19 \times 10^{-7}$ | $7.71 \times 10^{-6}$ |
| atherosclerosis DOID-1936 mouse GSE19286 sample 906 | 6/259 | $2.79 \times 10^{-7}$ | $1.55 \times 10^{-5}$ |
| Type 2 diabetes mellitus C0011860 mouse GSE2899 sample 45 | 6/268 | $3.41 \times 10^{-7}$ | $1.66 \times 10^{-5}$ |
| Hepatocellular carcinoma DOID-684 human GSE10393 sample 493 | 6/298 | $6.35 \times 10^{-7}$ | $2.48 \times 10^{-5}$ |
| Carcinoma, Hepatocellular C0019204 mouse GSE2127 sample 300 | 6/298 | $6.35 \times 10^{-7}$ | $2.48 \times 10^{-5}$ |

| Disease Perturbations from GEO up | | | |
|---|---|---|---|
| Term | Overlap | P-value | Adjusted P-value |
| intracranial aneurysm DOID-10941 human GSE26969 sample 744 | 13/163 | $2.01 \times 10^{-22}$ | $8.89 \times 10^{-20}$ |
| Hyperlipidemia C0020473 rat GSE3512 sample 38 | 12/277 | $2.24 \times 10^{-17}$ | $4.97 \times 10^{-15}$ |
| Hepatic Cirrhosis C0023890 rat GSE1843 sample 372 | 12/297 | $5.21 \times 10^{-17}$ | $7.70 \times 10^{-15}$ |
| Sepsis C0243026 rat GSE1781 sample 314 | 10/226 | $1.60 \times 10^{-14}$ | $1.77 \times 10^{-12}$ |
| NASH C0400966 human GSE24807 sample 185 | 10/275 | $1.15 \times 10^{-13}$ | $1.02 \times 10^{-11}$ |
| Burn C0006434 rat GSE802 sample 97 | 9/283 | $8.48 \times 10^{-12}$ | $6.26 \times 10^{-10}$ |
| Hepatic lipidosis C0015695 mouse GSE5538 sample 10 | 9/316 | $2.27 \times 10^{-11}$ | $1.44 \times 10^{-9}$ |
| Vitamin A Deficiency C0042842 rat GSE1600 sample 359 | 8/265 | $2.33 \times 10^{-10}$ | $1.29 \times 10^{-8}$ |
| Atherosclerosis C0004153 mouse GSE363 sample 193 | 8/299 | $6.06 \times 10^{-10}$ | $2.98 \times 10^{-8}$ |
| colitis DOID-0060180 mouse GSE34874 sample 800 | 8/345 | $1.87 \times 10^{-9}$ | $8.30 \times 10^{-8}$ |
